# Characterization of pyruvate dehydrogenase complex E1 alpha and beta subunits of *Mycoplasma synoviae*

**DOI:** 10.1101/551176

**Authors:** Shijun Bao, Xiaoqin Ding, Shengqing Yu, Chan Ding

**Author notes:** Corresponding author, (S J Bao), (C. Ding).

## Abstract

*Mycoplasma synoviae* (MS) is an important pathogen, causing enormous economic losses to the poultry industry worldwide every year. Therefore, the studies on MS will lay the foundation for diagnosis, prevention and treatment of MS infection. In this study, primers designed based on the sequences of pyruvate dehydrogenase complex (PDC) E1 alpha and beta subunit genes (*pdhA* and *pdhB*, respectively) of MS WVU1853 strain in GenBank were used to amplify the *pdhA* and *pdhB* genes of MS WVU1853 strain through PCR. Then the prokaryotic expression vectors pET-pdhA and pET-pdhB were constructed and were expressed in *Escherichia coli* BL21(DE3) cells. Subsequently, the recombinant proteins rMSPDHA and rMSPDHB were purified and anti-rMSPDHA and anti-rMSPDHB sera were prepared by immunizing rabbits, respectively. Finally, the subcellular localization of PDHA and PDHB in MS, binding activity of rMSPDHA and rMSPDHB to chicken plasminogen (Plg) and human fibronectin (Fn), complement-dependent mycoplasmacidal assays, and adherence and adherence inhibition assays were accomplished. The results showed that PDHA and PDHB were distributed both on the surface membrane and within soluble cytosolic fractions of MS cells. The rMSPDHA and rMSPDHB presented binding activity with chicken Plg and human Fn. The rabbit anti-rMSPDHA and anti-rMSPDHB sera had distinct mycoplasmacidal efficacy in the presence of guinea pig complement, and the adherence of MS to DF-1 cells pretreated with Plg was effectively inhibited by treatment with anti-rMSPDHA or anti-rMSPDHB sera. Hence, the study indicates that the surface-associated MSPDHA and MSPDHB are the adhesion-related factors of MS that contributes to bind to Plg/Fn and adhesion to DF-1 cells.

---

*Mycoplasma synoviae* (MS) is an important pathogen exerting significant economic impact on poultry industry worldwide. MS infections can lead to a range of diseases, from subclinical to severe ailments. In chickens, local infection of MS frequently causes subclinical disease of the upper respiratory tract (Buim et al., 2009; Khiari et al., 2010; Landman, 2014; Wetzel et al., 2010), while systemic infection of MS causes synovitis (Wetzel et al., 2010). Furthermore, coinfection of MS with other respiratory pathogens, such as Newcastle disease virus, infectious bursal disease virus, infectious bronchitis virus, *Escherichia coli*, and *Mycoplasma meleagridis*, can induce airsaculitis, thereby further increasing economic losses (Giambrone et al., 1977; Kleven, 1998; Limpavithayakul et al., 2016; May et al., 2007; Rhoades, 1977; Springer et al., 1974; Vardaman et al., 1975). In addition to respiratory tract tropism strains and arthropathic strains, oviduct tropism strains can induce eggshell apex abnormalities without any physical abnormalities (Limpavithayakul et al., 2016). Therefore, research on MS can establish the theoretical foundation for further development of vaccines, diagnostic reagents, and therapeutic drugs against MS infections.

The pyruvate dehydrogenase complex (PDC) catalyzes the oxidative decarboxylation of pyruvate to acetyl-CoA (Patel and Roche, 1990; Perham, 1991; Reed, 2001). Therefore, the PDC occupies a key position in the oxidation of glucose by linking the glycolytic pathway to the oxidative pathway of the tricarboxylic acid cycle. In prokaryotes and eukaryotes, the PDC consists of three catalytic enzymes: pyruvate dehydrogenase (E1), dihydrolipoamide acetyltransferase (E2), and dihydrolipoamide dehydrogenase (E3). The PDC E1(EC 1.2.4.1) is a thiamin diphosphate-dependent enzyme and catalyzes the oxidative decarboxylation of pyruvate, which is the rate-limiting step in the overall activity of the PDC (Linn et al., 1969a; Linn et al., 1969b; Patel et al., 2014). Previous studies have shown that the PDC E1 is a heterotetramers (α_2_β_2_) consisting of two alpha subunits and two beta subunits (Dahl et al., 1987; Patel et al., 2014; Payton et al., 1977). In some *Mycoplasma* spp., the E1 beta subunit of PDC acts as an immunogenic protein or plasminogen (Plg)/fibronectin (Fn) binding proteins (Dallo et al., 2002; Sun et al., 2014; Thomas et al., 2013); however, there are still no reports on PDC E1 of MS. Therefore, in the present study, the genes encoding the alpha and beta subunits of PDC of MS were amplified and expressed in *E. coli*, respectively, and their biological characteristics were investigated.

## MATERIALS AND METHODS

### Enzymes and main reagents

Restriction enzymes, T4 DNA ligase and PrimeSTAR^®^ HS DNA polymerase, were purchased from TaKaRa (Dalian, China). Complement sera (guinea pig source) were bought from the Control Institute of Veterinary Bioproducts and Pharmaceuticals (Beijing, China). The reagents used for cell culture were obtained from Gibco (Grand Island, NY, USA). Other chemicals used in this study were of analytical grade and purchased from Sigma-Aldrich (St. Louis, MO, USA) or Sangon Biotech (Shanghai, China).

### Bacterial strains, cell line, culture conditions, and plasmids

*M. synoviae* WVU1853 strain was obtained from the Chinese Veterinary Culture Collection Center (CVCC, Beijing, China) and cultured in Frey’s medium (Frey et al., 1968) at 37°C in an atmosphere of 5% CO_2_. *E. coli* strains DH5α and BL21(DE3) (TransGen, Beijing, China), which were used as prokaryotic host for cloning and expression, respectively, were cultured in Luria-Bertani (LB) broth or plated on LB agar supplemented with 50 μg/mL kanamycin at 37°C. DF-1 cells, a continuous cell line of chicken embryo fibroblasts, were obtained from the American Type Culture Collection (Manassas, VA, USA) and certified to be free of Mycoplasma contamination, and were grown in Dulbecco’s modified Eagle’s medium (DMEM) supplemented with 10% fetal bovine serum, 100 IU/mL penicillin, and 100 μg/mL streptomycin at 37°C in an atmosphere of 5% CO_2_. The pET-28a(+) expression vector was obtained from Novagen (Madison, WI, USA), and pET-InaZNEGFP was constructed in our previous study (Bao et al., 2015).

### Cloning and expression of pdhA and pdhB genes of MS, and protein purification

The complete genome sequence of MS strain WVU1853 available in GenBank (www.ncbi.nlm.nih.gov/genbank/) revealed that the open reading frame (ORF) of MS *pdhA* gene containing eight TGA codons encodes tryptophan in *Mycoplasma* spp., but acts as stop codons in *E. coli*. Therefore, primers pdhA-F1/pdhA-R1, pdhA-F2/pdhA-R2, pdhA-F3/pdhA-R3, pdhA-F4/pdhA-R4, pdhA-F5/pdhA-R5, pdhA-F6/pdhA-R6, pdhA-F7/pdhA-R7, pdhA-F8/pdhA-R8, and pdhA-F9/pdhA-R9 (Table 1) were designed for site-directed mutagenesis and used in overlapping PCRs to amplify *pdhA*, while primers pdhB-F/pdhB-R (Table 1) were employed to amplify *pdhB*. The full-length mutated *pdhA* and *pdhB* genes were amplified with the introduction of *Bam*HI and *Xho*I restriction enzyme sites, respectively, and the recombinant plasmids pET-pdhA and pET-pdhB were constructed and transformed into *E. coli* BL21(DE3) cells, respectively. Recombinant rMSPDHA and rMSPDHB were then expressed through 1.0 mM isopropyl β-D-1-thiogalactopyranoside (IPTG) induction and purified using Ni-NTA His-Bind^®^ Resin Kit (Novagen, San Diego, USA). The purified protein products were quantified by using BCA Protein Assay Kit (Thermo Scientific-Pierce, Rockford, IL, USA) and analyzed by 10% SDS-PAGE with Coomassie blue staining.

**Table 1.**
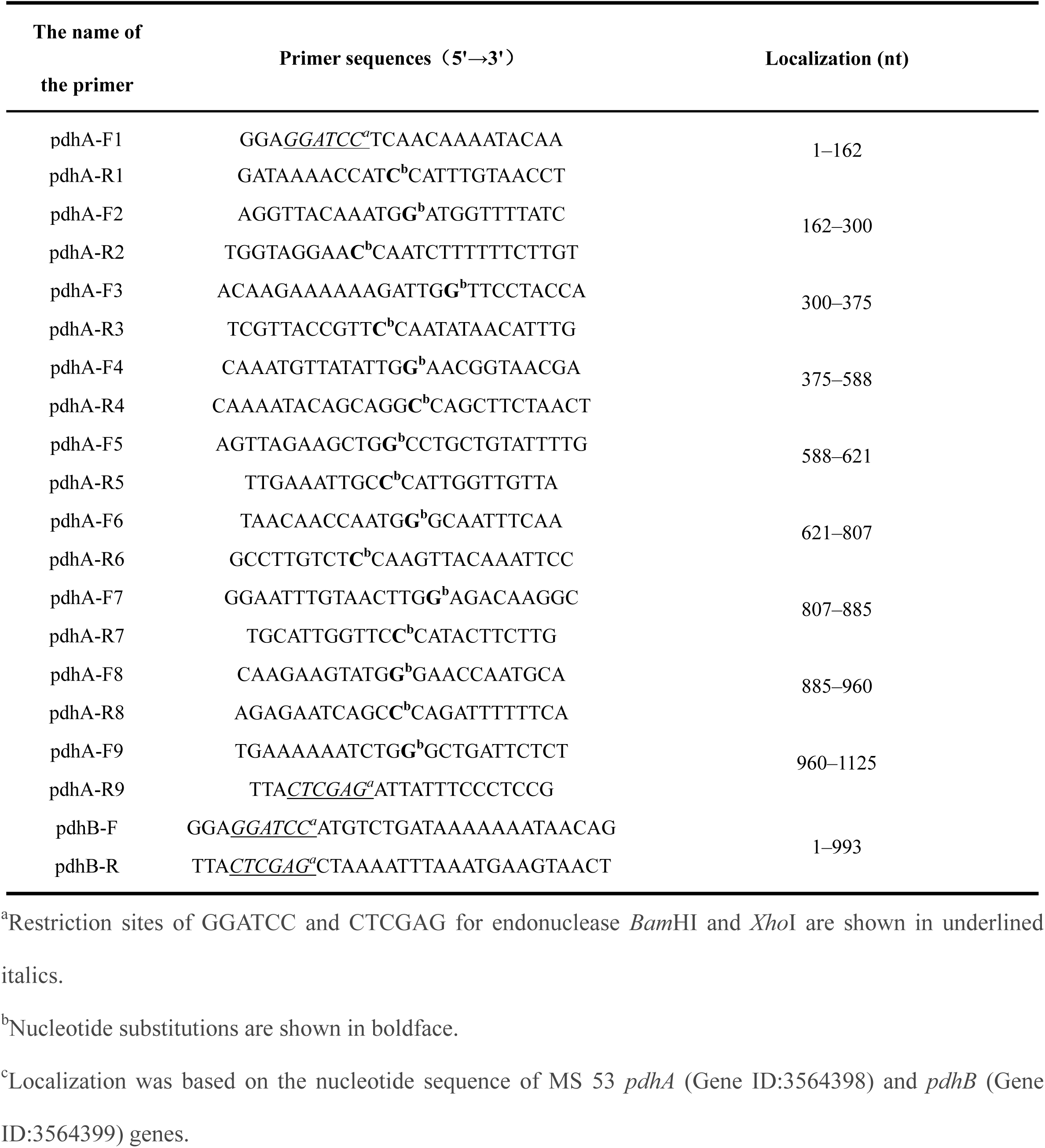
Primers used in this study.

### Preparation of anti-sera against rMSPDHA, rMSPDHB

The anti-sera against rMSPDHA, rMSPDHB, and MS whole cells were prepared by injecting female New Zealand White rabbits with rMSPDHA, rMSPDHB, and inactivated whole-cell MS, respectively. Three female rabbits were individually immunized four times by subcutaneous injection into the back with purified recombinant proteins (800 μg) or inactivated whole-cell MS (10^10^ CFU), mixed with Imject^®^ Alum adjuvant (Thermo Scientific-Pierce, Waltham, MA, USA), at 2-week intervals. Seven days after the fourth immunization, the rabbits were bled and the antibody titers were measured by indirect ELISA (Bao et al., 2014). Aliquots of the serum samples were the placed in 1.5-mL Eppendorf tubes and stored at −40°C for future use. All animal experiments were approved by the Experimental Animal Ethic and Welfare Committee of Gansu Agricultural University (China).

### Preparation of MS protein fractions

Membrane and cytosolic proteins fractions from MS were extracted using ReadyPrep™ Protein Extraction Kit (Membrane I) (Bio-Rad, CA, USA) according to the manufacturer’s instructions. Subsequently, the membrane and cytosolic proteins were dissolved in the same volume of protein solubilization buffer and equal volumes of the solutions were respectively subjected to ELISA or Western blot. The MS whole cell protein was prepared by using bacterial lysis buffer (Sangon, Shanghai, China) according to the manufacturer’s instructions. Protein quantitation was performed using BCA Protein Assay Kit (Beyotime Institute of Biotechnology, Nan Tong, China).

### Analysis of PDHA and PDHB distribution in MS whole cells

To confirm the distribution of PDHA and PDHB in MS whole cells, equal volumes of membrane and cytosolic proteins were subjected to western blot or ELISA. Western blot was performed as described previously (Bao et al., 2014). In brief, the gels were transferred onto nitrocellulose (NC) membranes (Whatman GmbH, Staufen, Germany) and blocked with 5% skim milk for 3 h at room temperature. Following three washes with PBST (3.2 mM Na_2_HPO_4_, 0.5 mM KH_2_PO_4_, 1.3 mM KCl, 135 mM NaCl, 0.05% Tween 20, pH 7.4), the NC membranes were incubated with rabbit anti-rMSPDHA or anti-rMSPDHB sera (dilution, 1:1000) at 4°C overnight. After washing thrice, the NC membranes were incubated with goat anti-rabbit IgG conjugated to horseradish peroxidase (HRP; Sigma-Aldrich; dilution, 1:8000) and the color reaction was examined using ECL Kit (Amersham Pharmacia Biotech, Piscataway, NJ, USA). Purified rMSPDHA and rMSPDHB (1.5 μg) were used as positive control and bovine serum albumin (BSA, 1.5 μg) was employed as negative control. The experiment was performed in triplicate and repeated three times.

For ELISA, the 96-well plates were coated with equal volumes of membrane and cytosolic proteins and incubated at 4°C overnight. After washing thrice with PBST, the wells were blocked with 5% skim milk in PBST at 37°C for 3 h. The plates were then washed thrice and rabbit anti-rMSPDHA sera or rabbit anti-rMSPDHB sera (100 μL/well; dilution, 1:1000) were added to the wells and incubated for 2 h at 37°C. After washing, goat anti-rabbit IgG-HRP was added to the wells (100 μL/well; Sigma-Aldrich; dilution, 1:5000) and the plates were incubated at 37°C for 1 h. Finally, the color reaction was conducted by adding soluble tetramethylbenzidine (TMB) substrate solution to the wells (100 μL/well; TIANGEN, Beijing, China) and incubating the plates for 10 min at room temperature. The reaction was stopped with 2 M H_2_SO_4_ addition. The absorbance was measured at A_450_ using a microplate reader (Bio-Tek Instruments, Winooski, USA). Purified rMSPDHA and rMSPDHB (10 μg/well) were used as positive control and BSA (10 μg/well) was employed as negative control. The experiment was performed in triplicate and repeated three times.

### Complement-dependent mycoplasmacidal assays

Complement-dependent mycoplasmacidal activity of rabbit anti-rMSPDHA or anti-rMSPDHB sera was determined as described previously (Bao et al., 2014). MS, grown to mid-logarithmic phase, was washed three times with PBS by centrifugation at 5,000×g for 10 min at 4°C, and re-suspended in PBS at the final concentration of 6×10^3^ CFU/mL. The reaction system was established as follows: 160 μL of MS suspension and 60 μL of rabbit anti-rMSPDHA or anti-rMSPDHB sera (1:5) were gently mixed in 1.5-mL Eppendorf tube and incubated at 37°C for 30 min. Then, 30 μL of diluted complement (1:10) were added, mixed and incubated at 37°C for 1 h. The reaction mixture (50 μL) was spread onto solid media in a 60-mm dish and incubated at 37°C in 5% CO_2_ for 7 days to count the colonies. Rabbit anti-MS sera and pre-immune rabbit sera were employed as positive and negative control, respectively. In addition, controls for complement and PBS were included. All sera (except for complement) used in the experiments were inactivated at 56°C for 30 min. Three independent experiments were performed in triplicate. The mycoplasmacidal coefficient was calculated as follows: [(CFU of pre-immune serum treatment - CFU of antiserum treatment)/(CFU of pre-immune serum treatment)] × 100.

### Binding activity of rMSPDHA and rMSPDHB to Plg and Fn

Western blot and ELISA were used to determine the binding activity of rMSPDHA and rMSPDHB to chicken Plg (Cell Sciences, MA, USA) and human Fn (Sigma-Aldrich) as described previously (Bao et al., 2014). In brief, MS rMSPDHA or rMSPDHB was transferred onto nitrocellulose membranes, incubated with chicken Plg or human Fn, and blots were developed with ECL Kit (Amersham Pharmacia Biotech) according to the manufacturer’s instructions.

ELISA was performed to verify the ability of the proteins to bind to Plg and Fn. In brief, the wells of the ELISA plate were coated with chicken Plg or human Fn at a concentration of 50 ng/well and incubated at 4°C overnight. After washing, a range of concentrations of rMSPDHA or rMSPDHB (0, 1, 5, 10, 15, 20, 25, or 30 μg/mL in PBST) were added to the wells and incubated for 2 h at 37°C. Then, the plates were incubated with rabbit anti-rMSPDHA or anti-rMSPDHB sera (100 μL/well; dilution, 1:1000) at 37°C for 1.5 h. After washing, goat anti-rabbit IgG-HRP was added to the wells (100 μL/well; Sigma-Aldrich; dilution, 1:5000), and the plates were incubated at 37°C for 1 h. Subsequently, the color reaction was performed and the absorbance was measured at A_450_ using a microplate reader (Bio-Tek Instruments). BSA were used as negative control and all experiments were performed in triplicate and repeated thrice.

### Adherence and adherence inhibition test

Adherence and adherence inhibition assays were performed as previously described (Bao et al., 2015). *E. coli* surface display vectors pET-InaZN-EGFP-pdhA/pET-InaZN-EGFP-pdhB were constructed, and *E.coli* BL21 (DE3) cells harbouring InaZN-EGFP-pdhA or InaZN-EGFP-pdhB fusion protein were induced by IPTG, centrifuged at 2500 g for 10 min and washed three times with DMEM. Then the DF-1 cells grown to monolayers in 35-mm dishes were washed with DMEM and the pdhA-positive or pdhB-positive *E. coli* cells were added to DF-1 cells at 100 multiplicity of infection (MOI) for 2h at 37°C in 5% CO2. After washing with PBS to remove non-adherent *E. coli* cells, 1 ml of 4% paraformaldehyde was added to the monolayers, and the plate was then incubated at room temperature for 10 min. After washing five times with PBS, the cell membranes were labelled with 500 µl of 10 µmol l^-1^ 1.1′-dioctadecyl-3,3,3′,3′-tetramethylindocarbocyanine perchlorate (Beyotime Institute of Biotechnology, Jiangsu, China) at room temperature for 10 min. After washing five additional times with PBS, the cell nuclei were labelled with 500 µl of 0.1 µg ml^-1^ of 4′, 6-diamidino-2-phenylindole (Beyotime Institute of Biotechnology) at room temperature for 10 min and then washed three times with PBS. Finally, the cells were mounted on slides and observed by fluorescent microscopy to evaluate the adhesion properties of the cells. The induced *E. coli* BL21(DE3) cells harboring InaZN-EGFP fusions acted as blank control, and all the experiments were performed in triplicate and repeated thrice.

To further validate the adherence function of MSPDHA and MSPDHB, adherence/adherence inhibition assay was conducted based on a bacteriological assay as previously described (Chen et al., 2011; Song et al., 2012) with slight modifications. For the adherence assay, DF-1 cells grown to a monolayer in a 35-mm dish were washed thrice with PBS and incubated with Plg (10 μg/mL) for 2 h at 37°C. The cells were then washed with DMEM and the MS WVU1853 strain was added at a multiplicity of infection of 200 and incubated for 2 h at 37°C and 5% CO_2_. Non-adherent MS cells were removed by washing and the infected cells were lysed with 0.25% trypsin (Gibco) and serial dilutions of the cell lysate were then respectively plated onto solid medium and incubated for 96 h at 37°C and 5% CO_2_. The MS colonies were counted to determine the adherence frequency. For adherence inhibition assay, the MS WVU1853 strain was incubated with rabbit anti-rMSPDHA/anti-rMSPDHB sera or rabbit anti MS positive sera for 1 h at 37°C and then used to infect the DF-1 cells as described earlier. The pre-immune rabbit sera acted as negative control and all experiments were performed in triplicate and repeated thrice. The percent inhibition was calculated using the following formula: (CFU of negative control - CFU of anti-serum treatment)/(CFU of negative control)] × 100.

### Statistical analysis

All data are expressed as the mean ± standard deviation of n independent measurements. The statistical significance of the intergroup differences was evaluated using Student’s t-test. The SPSS software (SPSS, Chicago, IL, USA) was used to calculate and analyze and the level of significance was set at P < 0.05 or P < 0.01.

## RESULTS

### Expression of MS pdhA and pdhB genes, and purification of recombinant proteins

The full-length mutated *pdhA* and *pdhB* genes were amplified with the designed primers using overlapping PCR, and the recombinant plasmid pET-pdhA and pET-pdhB were constructed. Sequence analysis indicated that the *pdhA* and *pdhB* genes were 1125 and 993 bp in length, respectively. The tryptophan codons (TGA) in the *pdhA* gene were successfully mutated to TGG. Theoretically, *pdhA* gene encodes a 374-amino-acid protein with a relative molecular weight of approximately 40.642 kDa, while *pdhB* gene encodes a 330-amino-acid protein with a relative molecular weight of approximately 35.854 kDa.

The recombinant plasmids of pET-pdhA and pET-pdhB vectors were respectively transformed into *E. coli* BL21(DE3) cells. The obtained recombinant proteins rMSPDHA and rMSPDHB were expressed by 1.0 mM IPTG induction. The SDS-PAGE results showed that the proportions of recombinant fusion proteins, rMSPDHA and rMSPDHB, were higher in the supernatant than those in the sediment, and their apparent molecular weights were approximately 44 and 39 kDa, respectively (Fig. 1). The purified recombinant proteins presented a single band,respectively (Fig. 2).

**Fig. 1.**
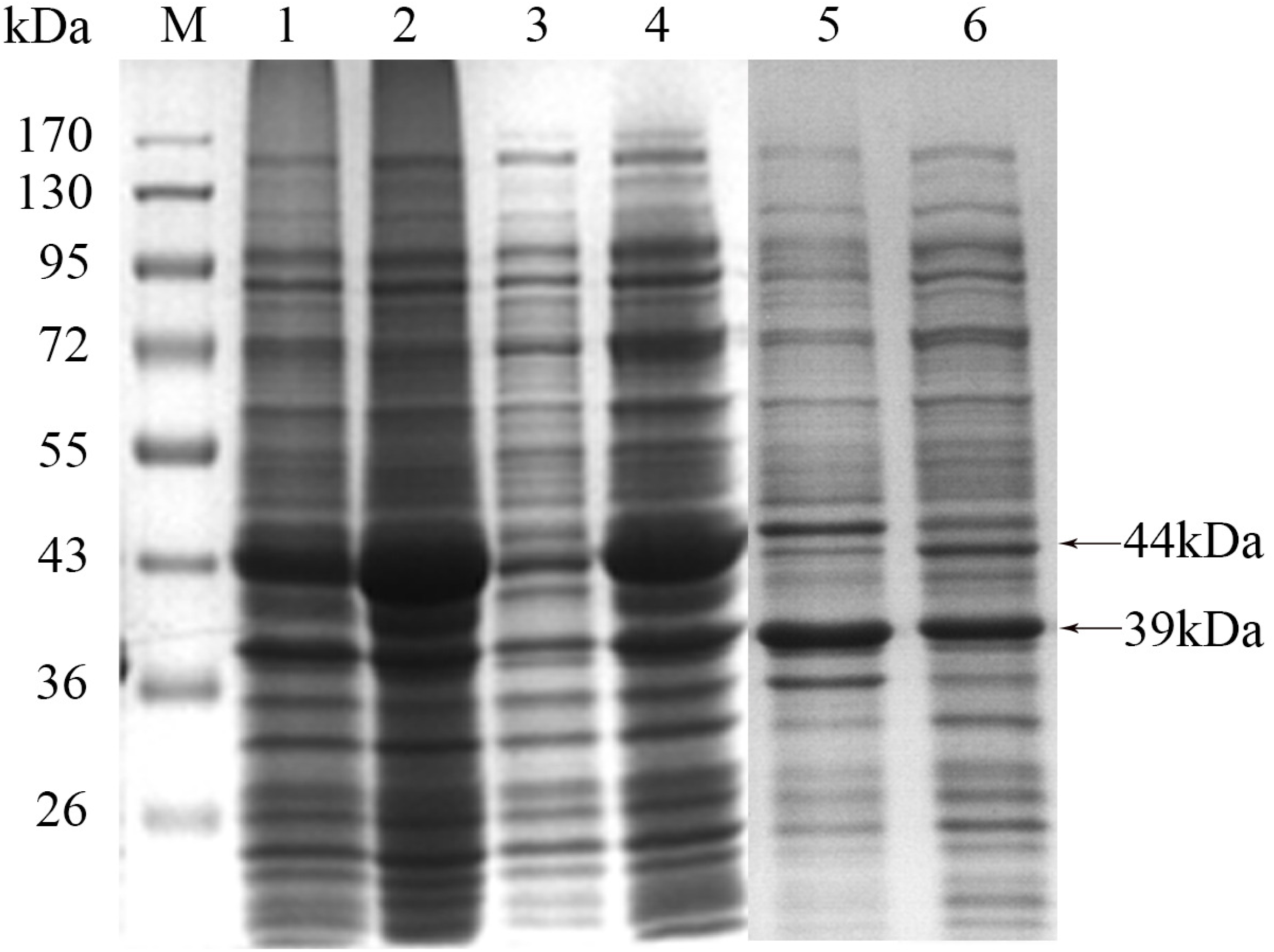
Analysis of the expression of recombinant proteins using SDS-PAGE followed by Coomassie blue staining. M: Pre-stained protein molecular weight marker (SM1811, Fermentas); Lane 1: Precipitation of lysate of *E. coli* BL21(DE3) cells transformed by pET-pdhA; Lane 2: Supernatant of lysate of *E. coli* BL21(DE3) cells transformed by pET-pdhA; Lane 3: Total cellular proteins of *E. coli* BL21 (DE3) cells transformed by pET-28a (+); Lane 4: Total cellular proteins of *E. coli* BL21 (DE3) cells transformed by pET-pdhA; Lane 5: Precipitation of lysate of *E. coli* BL21(DE3) cells transformed by pET-pdhB; Lane 6: Supernatant of lysate of *E. coli* BL21(DE3) cells transformed by pET-pdhB.

**Fig.2.**
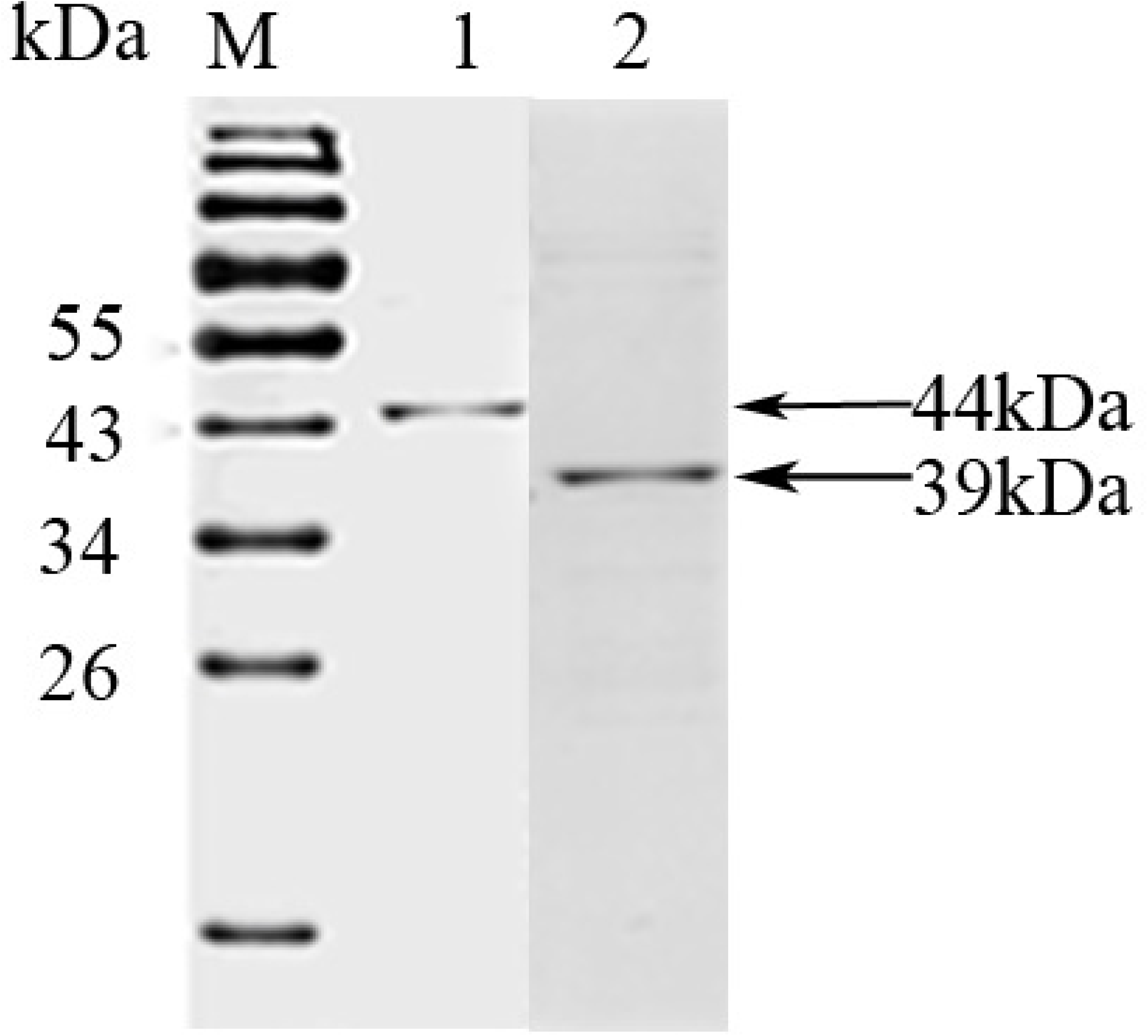
Analysis of purified recombinant proteins using SDS-PAGE followed by Coomassie blue staining. M: Pre-stained protein molecular weight marker (SM1811, Fermentas); Lane 1: Purified rMSPDHA; Lane 2: Purified rMSPDHB.

### Preparation of rabbit anti-sera against rMSPDHA, rMSPDHB, and MS whole cells

One week after fourth immunization, the rabbits were bled and the reactivity and specificity of the rabbit antisera were respectively tested by ELISA and western blot. The results showed that the ELISA antibody titers of the anti-rMSPDHA, anti-rMSPDHB, or anti MS whole cells sera were higher than 1:20,000. Western blot results demonstrated that both anti-rMSPDHA and anti-rMSPDHB sera can specifically combine with MS bacterial protein and purified rMSPDHA or rMSPDHB (Fig. 3).

**Fig. 3.**
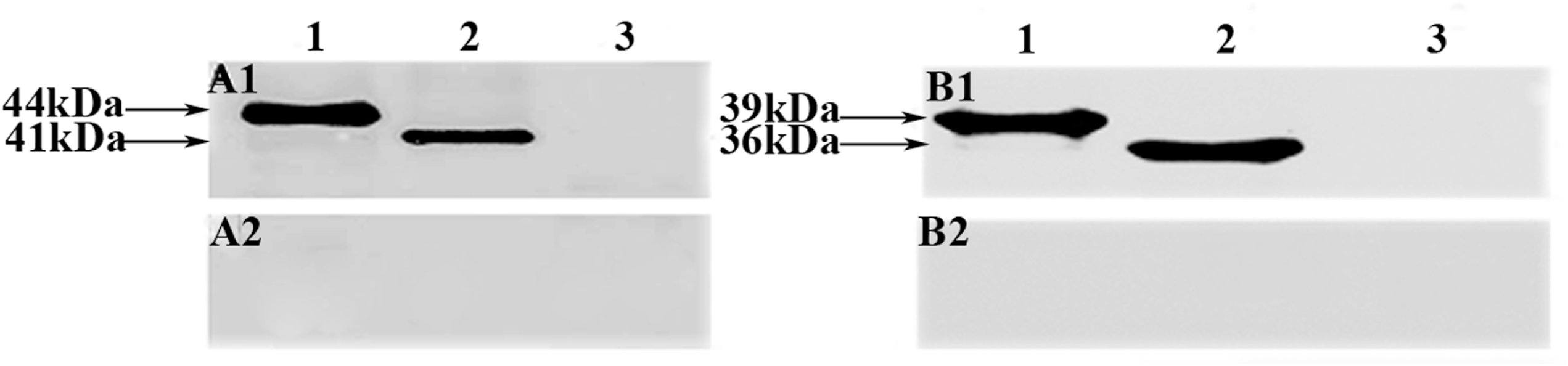
Western blot analysis of rabbit anti-rMSPDHA and anti-rMSPDHB sera. A1: Western blot analysis of rabbit anti-rMSPDHA serum; A2: Negative serum from rabbit; Lane 1: Purified rMSPDHA protein; Lane 2: Total proteins of MS WVU1853 strain; Lane 3: BSA control. B1: Western blot analysis of rabbit anti-rMSPDHB serum; B2: Negative serum from rabbit; Lane 1: Purified rMSPDHB protein; Lane 2: Total proteins of MS WVU1853 strain; Lane 3: BSA control

### Localization of MSPDHA and MSPDHB

MS PDHA and MS PDHB were detected in the cell membrane fraction proteins (Fig. 4; Lanes 2, 6) and the cell soluble cytosolic fraction proteins of MS (Fig. 4; Lanes 3, 7). Purified rMSPDHA, rMSPDHB (Fig. 4; Lanes 1, 5), and BSA (Fig. 4; Lanes 4, 8) were employed as positive and negative controls, respectively. The anti-rMSPDHA and anti-rMSPDHB sera could specifically combine with the protein of approximately 41kDa and 36 kDa, respectively, while the band size of the binding recombinant protein was about 44 kDa and 39 kDa, respectively. However, both anti-rMSPDHA and anti-rMSPDHB sera had no binding band with BSA, suggesting that MS PDHA and PDHB were present in both the membrane and soluble cytosolic protein fractions of MS cells. And the results also indicated that the content of MS PDHA and PDHB in the membrane fractions was higher than that in the cytosolic fractions. ELISA results (Fig. 5) also revealed that the content of MS PDHA and PDHB in the membrane fractions was higher than that in the cytosolic fractions (P<0.01).

**Fig. 4.**
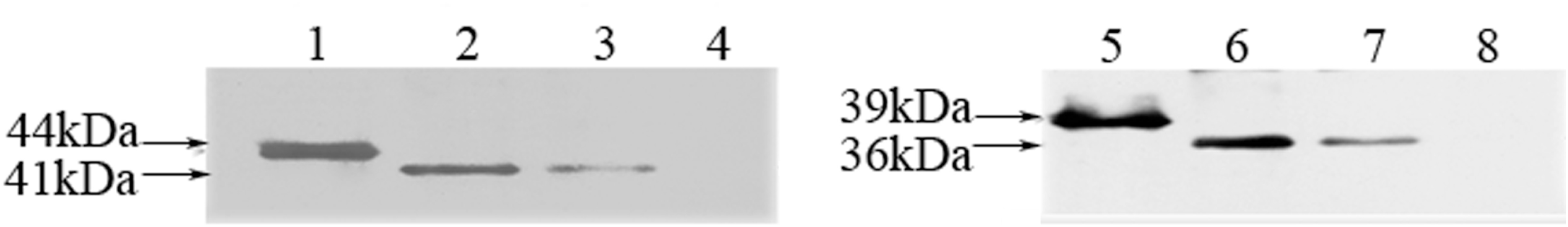
Western blot analysis of localization of MSPDHA and MSPDHB. Panel A: Western blot analysis using anti-rMSPDHA; Panel B: Western blot analysis using anti-rMSPDHB; Lanes 1, 5: Purified rMSPDHA and rMSPDHB, used as positive control; Lanes 2, 6: Membrane proteins of MS; Lanes 3, 7: Cytosolic proteins of MS; Lanes 4, 8: BSA was used as negative control.

**Fig. 5.**
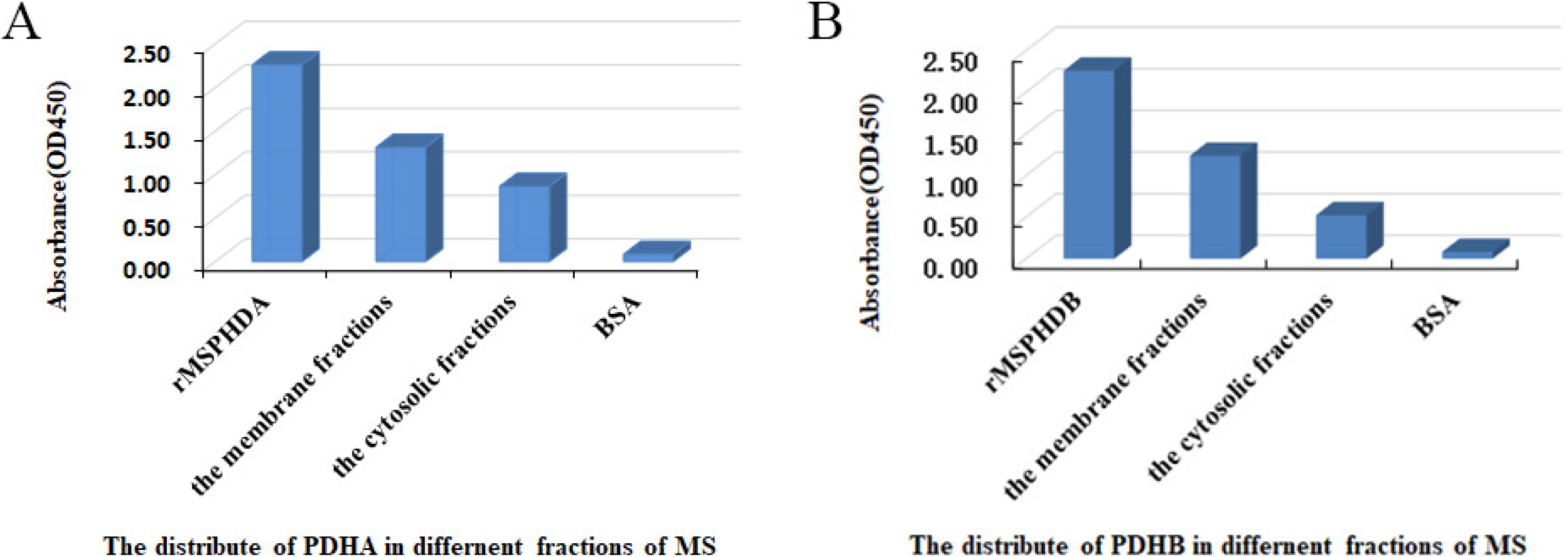
ELISA of MSPDHA and MSPDHB localization. Panel A: ELISA of PDHA distribution in the membrane and soluble cytosolic protein fractions of MS cells using anti-rMSPDHA; Panel B: ELISA of PDHB distribution in the membrane and soluble cytosolic protein fractions of MS cells using anti-rMSPDHB.

### Complement-dependent mycoplasmacidal assays

As shown in Table 2, Complement-dependent mycoplasmacidal assays revealed obvious difference in mycoplasmacidal activity between anti-rMSPDHA/anti-rMSPDHB sera and pre-immune rabbit serum (P<0.01). In addition, the mycoplasmacidal activity of the rabbit anti-rMSPDHA was more effective than that the anti-rMSPDHB sera, and the mycoplasmacidal rates were 65.6% and 29.89%, respectively.

**Table 2.** Mycoplasmacidal activity of rabbit anti-rMSPDHA/anti-rMSPDHB sera

| Sera Type | Mean CFU of MS ($10^4$) | Mycoplasmacidal coefficient (%) |
| --- | --- | --- |
| MS positive sera | 10 | 96.5* |
| Rabbit anti-rMSPDHA sera | 100 | 65.6* |
| Rabbit anti-rMSPDHB sera | 204 | 29.89* |
| pre-immune rabbit sera | 291 | — |

\*P< 0.01, compared with the corresponding group using pre-immune rabbit serum.

### Binding activity of rMSPDHA and rMSPDHB to Plg and Fn

The ability of rMSPDHA and rMSPDHB to bind to Plg and Fn was examined by western blot and ELISA. The western blot analyses showed that both chicken Plg and human Fn bind to rMSPDHA and rMSPDHB, as indicated by unique bands at 44 and 39 kDa, respectively (Fig. 6). ELISA revealed that both rMSPDHA and rMSPDHB could bind to immobilized chicken Plg or human Fn in a dose-dependent manner (Fig. 7). However, BSA, which was employed as the negative control, showed no obvious binding effect with either chicken Plg or human Fn (Fig. 7).

**Fig.6.**
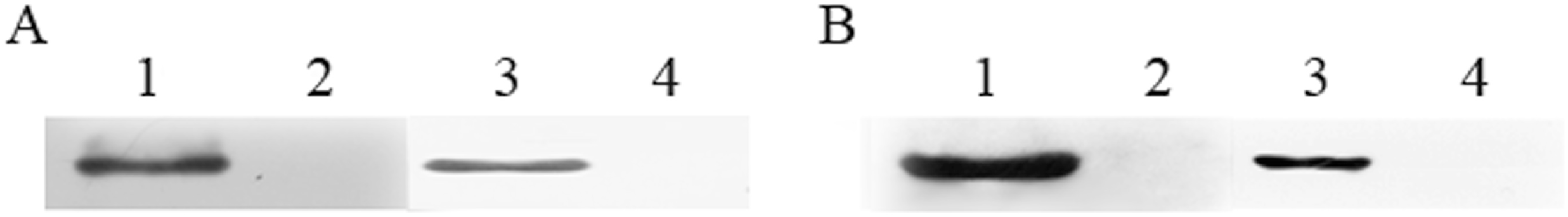
Western blot analysis of the binding ability of rMSPDHA and rMSPDHB to chicken Plg or human Fn. Panel A: Western blot analysis using anti-rMSPDHA; Panel B: Western blot analysis using anti-rMSPDHB; Lane 1: Chicken Plg; Lane 3: Human Fn; Lanes 2, 4: BSA.

**Fig.7.**
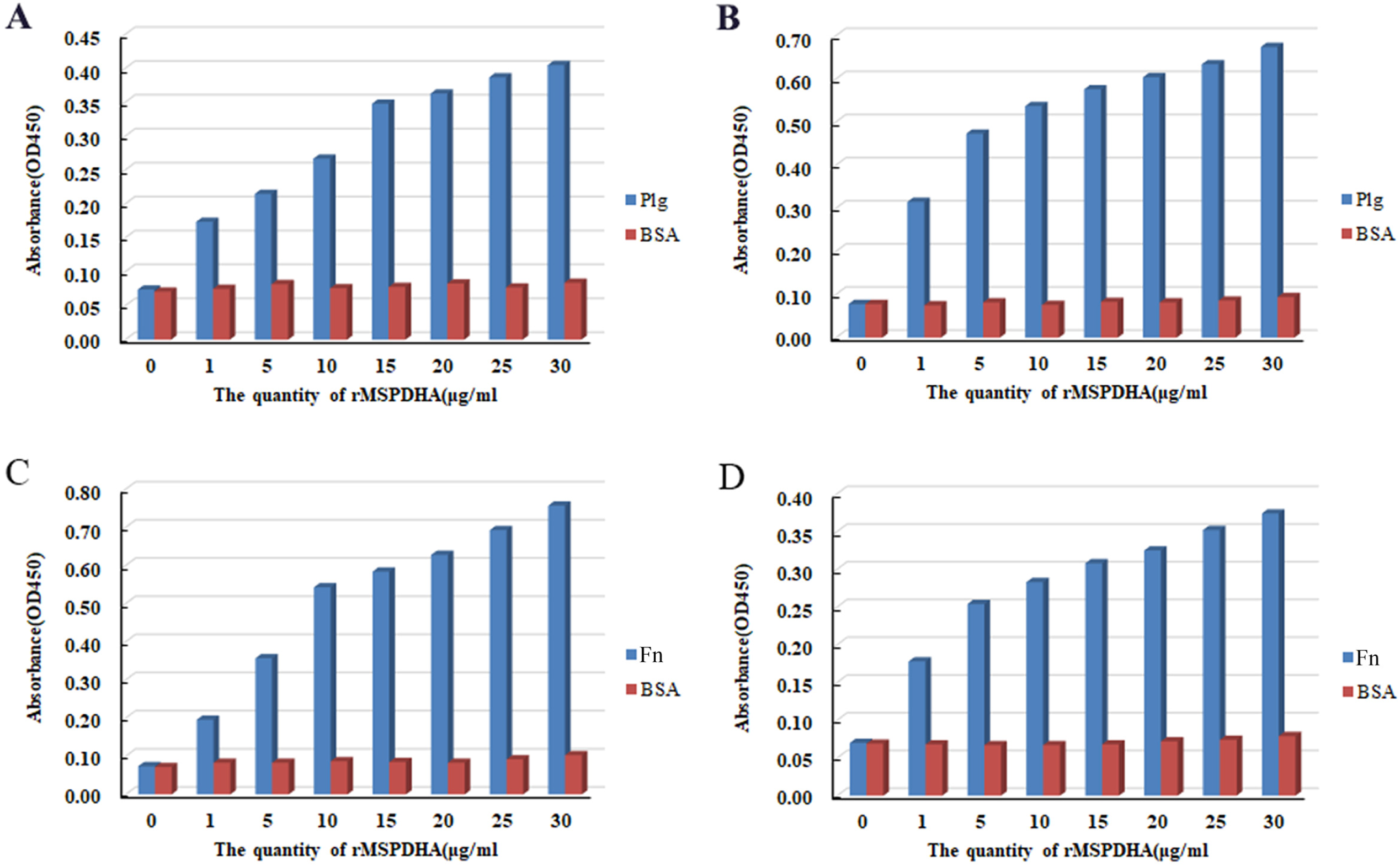
ELISA of MSPDHA and MSPDHB binding to chicken Plg or human Fn. (A) ELISA of the binding ability of rMSPDHA to chicken Plg. (B) ELISA of the binding ability of rMSPDHB to chicken Plg. (C) ELISA of the binding ability of rMSPDHA to human Fn. (D) ELISA of the binding ability of rMSPDHB to human Fn.

### Adherence and adherence inhibition assays

Adherence and adherence inhibition tests of *E. coli* cells harboring InaZNEGFP-PDHA or InaZNEGFP-PDHB to DF-1 cells showed that the PDHA-and PDHB-positive *E. coli* transformants were able to specifically attach to DF-1 cells, which can be inhibited by rabbit anti-rMSPDHA/anti-rMSPDHB sera (Fig. 8). However, the induced *E. coli* transformants harboring pET-InaZNEGFP were unable to attach to DF-1 cells (Fig. 8), indicating that both MS PDHA and MS PDHB are adhesion-related factors.

**Fig.8.**
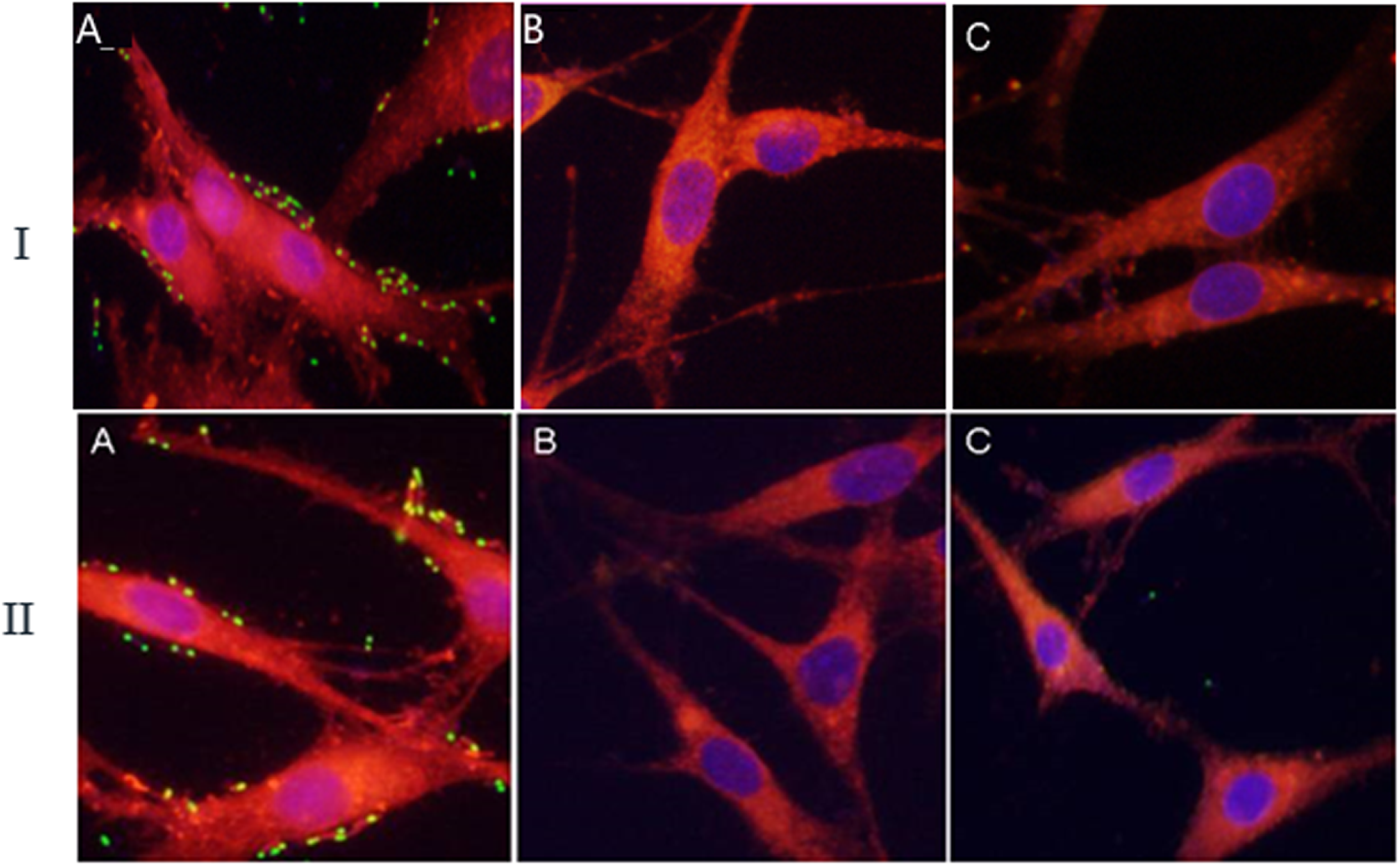
Adherence and adherence inhibition assays. IA: Adherence of *E. coli* cells harboring InaZNEGFP-PDHA to DF-1 cells; IB: Adherence of *E. coli* cells harboring InaZNEGFP to DF-1 cells; IC: Adherence inhibition by rabbit anti-rMSPDHA sera. IIA: Adherence of *E. coli* cells harboring InaZNEGFP-PDHB to DF-1 cells; IIB: Adherence of *E. coli* cells harboring InaZNEGFP to DF-1 cells; IIC: Adherence inhibition by rabbit anti-rMSPDHB sera.

To further validate the effect of MSPDHA and MSPDHB on the adherence of MS to DF-1 cells, the adherence and adherence inhibition values were calculated. As table 3 shows that the rabbit anti-rMSPDHA/anti-rMSPDHB sera had significant inhibitory effect on the adherence of MS to DF-1 cells (P< 0.01). There was apparent difference in adherence inhibition rate between the anti-rMSPDHA/anti-rMSPDHB sera and pre-immune rabbit serum (P< 0.01), and the anti-rMSPDHA sera were more effective than the anti-rMSPDHB sera at inhibiting adherence (P< 0.01). Thus, these results demonstrated that MSPDHA and MSPDHB are adhesion-related proteins.

**Table 3.** CFU values of different treatment groups.

| Serum | Mean CFU of MS($10^4$) | Adhesion inhibition rate(%) |
| --- | --- | --- |
| Rabbit anti MS positive sera | 60 | 93.33* |
| Rabbit anti-rMSPDHA sera | 132 | 85.33* |
| Rabbit anti-rMSPDHB sera | 170 | 81.11* |
| pre-immune rabbit sera | 900 | —— |

\*P < 0.01, compared with the corresponding group using pre-immune rabbit serum(negative control).

## DISCUSSION

MS is a major pathogen of avian, which is widely distributed around the world and leads to serious economic losses in the world every year. Therefore, the MS association studies will lay the foundation for diagnosis, prevention and treatment of MS infection. Prior to this study, some major immunogenic proteins of MS isolates were identified, including PDC E1 alpha and beta subunit, elongation factor Tu (EF-Tu), enolase, etc. (Bao et al., 2017; Bercic et al., 2008). As a component of the PDC, the PDC E1 not only plays the important role in energy metabolism, but performs additional biological functions, including immunogenicity, adhesion and invasion of pathogenic microorganisms to host cells(Sun et al., 2014; Vastano et al., 2014). Currently, PDC E1 from various plants and yeasts have been separated and identified, but there have been only a few reports on PDC E1 from bacteria. Bacteria-derived PDC E1 catalyzes ethanol production through Entner-Doudoroff pathway, rather than pyruvate generation through glycolysis. Therefore, studies on PDC E1, especially those on PDC E1 from thermophilic bacteria (Van Zyl et al., 2014), have been mainly focused on its biocatalysis, because it is safe, pollution-free, and highly efficient (Taylor et al., 2009). Nevertheless, no study on the PDC E1 of MS has been reported up to present.

In this study, the pdhA and pdhB of MS were amplified, optimized and expressed in a prokaryotic system, respectively. And both recombinant proteins could express in BL21(DE3) in soluble form. Then anti-rMSPDHA/anti-rMSPDHB sera were prepared. These laid the foundation for study on other biological functions of MS PDHA and PDHB. Subsequently, the immunogenic proteins with good immunogenicity of MSPDHA and MSPDHB were confirmed by indirect ELISA and western blot, consistent with previous research(Bao et al., 2017). The similar finding has been made in *M. bovis* (Sun et al., 2014).

That the MSPDHA and MSPDHB are present in both the soluble cytosolic fraction and the membrane fraction of MS has been demonstrated, and with higher distribution in the cell membrane fraction than in the cytosolic fraction (Fig.4 and 5). This analysis demonstrates that MSPDHA and MSPDHB are respectively the membrane-associated protein in MS, suggesting that MSPDHA and MSPDHB might possess various biological functions besides the catalytic activity catalyst, such as some membrane-associated proteins have been found in previous studies (Bao et al., 2014; Gao et al., 2018; Salzillo et al., 2017). We suggest that the distribution in the cell membrane fraction of MSPDHA and MSPDHB is consistent with the etiology of adherence to host cells.

We have also demonstrated that rabbit anti-rMSPDHA/anti-rMSPDHB sera displayed a significant complement-dependent mycoplasmacidal effect, and the mycoplasmacidal activity of the rabbit anti-rMSPDHA was more effective than that the anti-rMSPDHB sera (Table 2), the similar biological function of other proteins of MS has been previously reported(Bao et al., 2014; Gao et al., 2018). This suggested that MSPDHA and MSPDHB should play an important role in host immunity against MS infection.

The binding activity of MSPDHA and MSPDHB to chicken Plg and human Fn has been confirmed in the present study. The western blot analyses demonstrate the binding specificity of MSPDHA and MSPDHB to chicken Plg and human Fn (Fig. 6), and the ELISA revealed that the binding of MSPDHA and MSPDHB to chicken Plg or human Fn in a dose-dependent manner. The PDHB acts as fibronectin and plasminogen-binding protein has already been demonstrated in other microorganisms (Dallo et al., 2002; Salzillo et al., 2017; Thomas et al., 2013), but the binding activity of MSPDHA to chicken Plg and human Fn was first confirmation in this study. Those suggested that MSPDHA and MSPDHB are the major Plg/Fn-binding protein in MS; this could be of great importance for Mycoplasma establishment in the host. Therefore, we speculate that a-enolase is involved in MS adhesion to DF-1 cells.

To confirm the adhesion activity of MSPDHA and MSPDHB to host cells, both E. coli cells harboring InaZNEGFP-pdhA/InaZNEGFP-pdhB and MS were subjected to the adherence and inhibition assay in vitro. In this work the adhesion ability of MSPDHA and MSPDHB to DF-1 cells has been demonstrated, and this adhesion was effectively inhibited by the addition of anti-rMSPDHA/anti-rMSPDHB sera, suggesting that MSPDHA and MSPDHB are adhesion-related proteins on MS cell membrane surface. Based on this, we speculate that MSPDHA and MSPDHB may participate in adhesion, colonization, and invasion of MS to host cells, and may be involved in various clinical and pathologic sequelae of MS infection, such as synovitis, tenosynovitis and arthritis.

## CONCLUSIONS

The studies show that the MSPDHA and MSPDHB are surface-exposed proteins with excellent immunogenicity, and that rabbit anti-rMSPDHA/anti-rMSPDHB sera had a sinificant complement-dependent mycoplasmacidal effect. Furthermore, the findings also demonstrated the binding ability of MSPDHA and MSPDHB to chicken Plg and human Fn, as well as the adherence to DF-1 cells. These results suggested that MSPDHA and MSPDHB play important role in MS metabolism, infection, and immunity, providing the basis for further research on the functions of MSPDHA and MSPDHB.

## ACKNOWLEDGMENTS

We are thankful to the Laboratory of Veterinary Lemology, Gansu Agricultural University, China, and the Key Open Laboratory of Shanghai Veterinary Research Institute, CAAS, for providing key laboratory equipment.

